# Validating the genus *Pocheina* (Acrasidae, Heterolobosea, Excavata) leads to the recognition of three major lineages within Acrasidae

**DOI:** 10.1101/2024.10.05.616810

**Authors:** Alexander K. Tice, Kevin Regis, Timothy E. Shutt, Frederick W. Spiegel, Matthew W. Brown, Jeffery D. Silberman

## Abstract

*Pocheina* and *Acrasis* are two genera of heterolobosean sorocarpic amoebae within Acrasidae that have historically been considered close relatives. The two genera were differentiated based on their differing fruiting body morphologies. The validity of this taxonomic distinction was challenged when a SSU rRNA phylogenetic study placed an isolate morphologically identified as ‘*Pocheina*’ *rosea* within a clade of *Acrasis rosea* isolates. The authors speculated that pocheinoid fruiting body morphology might be the result of aberrant *A. rosea* fruiting body development, which if true, would nullify this taxonomic distinction between genera. To clarify Acrasidae systematics, we analyzed SSU rRNA and ITS region sequences from multiple isolates of *Pocheina, Acrasis*, and *Allovahlkampfia* generated by PCR and transcriptomics. We demonstrate that the initial SSU sequence attributed to ‘*P. rosea*’ originated from an *A. rosea* DNA contamination in its amplification reaction. Our analyses, based on morphology, SSU and 5.8S rRNA genes phylogenies, as well as comparative analyses of ITS1 and ITS2 sequences, resolve Acrasidae into three major lineages; *Allovahlkampfia* and the strongly supported clades comprising *Pocheina* and *Acrasis*. We confirm that the latter two genera can be identified by their fruiting body morphologies.

## INTRODUCTION

In 1873, Cienkowski described a microorganism he found on collections of dead lichenized wood in Russia (Cienkowski, 1873). Its fruiting body (sorocarp) was pink in color with a stalk consisting of a row of wedged shaped cells supporting a globular mass of spores at its apex. Each spore was said to contain pinkish cytoplasm and a nucleus, and when spores germinated a limax shaped amoeba with pink cytoplasm emerged. Cienkowski’s description of ‘*Guttulina rosea’* was the first of a non-dictyostelid sorocarpic amoeba (cellular slime mold) (Cienkowski, 1873). Aside from transferring the organism to the newly erected genus *Pocheina* (due to the recognition that the genus name *Guttulina* was already in use; Loeblich and Tappan, 1961), no work was done on the organism until its rediscovery in the 1970’s (Raper, 1973). A second species of *Pocheina* was later described, *P. flagellata;* named because anteriorly biflagellated cells as well as limax shaped amoebae emerged upon spore germination (Olive et al., 1983).

Another sorocarpic amoeba was later discovered by Van Teighem (1880), named *Acrasis granulata*. It was found fruiting on spent beer yeast as columnar rows of spores, brownish in color, that hatched amoeboid cells. Eighty years later, Olive & Stoianovitch (1960) named a new species to the genus *Acrasis, Ac. rosea*, because it matched the unillustrated text description of *Ac. granulata. Acrasis rosea* was found fruiting on collections of leaves and inflorescences of *Phragmites* sp. grass, and its spores germinated to produce limax shaped amoebae with pinkish-orange cytoplasm (Olive and Stoianovitch, 1960). The fruiting bodies of *Acrasis* differed from those of *Pocheina* in that they formed chains of spores rather than a globose mass at the apex of the stalk cells (Olive and Stoianovitch, 1960).

Olive et al. (1983) first proposed that *Pocheina* and *Acrasis* were closely related. Subsequently, they were placed with the vahlkampfiid amoebae into Heterolobosea (Page and Blanton 1985) because of the eruptive motion of the pseudopodia during locomotion of the amoeboid trophic cells, similarities in mitochondrial cristae structure (flattened discoidal cristae), and the close association of the mitochondria and endoplasmic reticulum (Dykstra, 1977; Olive, 1975; Page and Blanton, 1985; Pánek et al., 2016). The exact relationship between *Acrasis* and *Pocheina* remained unclear. Despite these morphological and ultrastructural similarities, *Pocheina* and *Acrasis* were always maintained as separate genera based primarily on sorocarp morphology (Dykstra, 1977; Page and Blanton, 1985).

In the first molecular phylogenetic study to include numerous geographically distributed isolates of ‘*Ac. rosea’*, it was shown that what was once thought to be merely morphological plasticity in the fruiting bodies among different isolates, were phylogenetically significant characteristics that could be used in conjunctions with molecular data to delineate species (Brown et al., 2012). Based on the congruence of morphology and molecular phylogenetic data using the nuclear encoded SSU rRNA gene (SSU) sequence, at least four distinct species of *Acrasis* exist (Brown et al., 2012). Included in this study was a partial SSU sequence generated from uncultured fruiting bodies, each topped with a globular spore mass, picked directly from its natural substrate, i.e., the morphotype typical of *Pocheina*. Surprisingly, this putative *Pocheina* (“*P. rosea”*) sequence was nested in a clade that contained all verified isolates of *A. rosea* (Brown et al., 2012). This led the authors to suggest that slight alterations during the development of *Ac. rosea* may be responsible for the formation of the chainless sorocarps that have previously been identified as *Pocheina* (Brown et al., 2012). This hypothesis was supported by the observation that long-term cultured isolates of *Ac. rosea* and *Ac. helenhemmesae* occasionally produce sorocarps with a globose spore mass atop a cellular stalk (Brown et al., 2010, 2012). If true, then the morphological difference ascribed to the fruiting bodies of *Acrasis*, and especially *Pocheina* would be taxonomically uninformative. Although the phylogenetic results were interpreted as best we could with the available data, for a variety of reasons, we remained suspicious of the ‘*Pocheina*’ isolate’s position within *Acrasis* for the following reasons. Slight variations in sorocarp morphology among species of *Acrasis* were representative of a large amount of molecular divergence in the SSU rRNA gene sequence among the different species. Additionally, the sorocarp morphology in previous cultures of the two known species of *Pocheina*, (*P. rosea* and *P. flagellata*; Cienkowski, 1873; Olive et al., 1983; Raper, 1973) remained stable through passaging. No culture of either species of *Pocheina* has been known to produce sorocarps that resemble sorocarps of any of the known species of *Acrasis* (Olive and Stoianovitch, 1960; Olive et al., 1983; Raper, 1973). Thus, the position of ‘*P. rosea*’ in our SSU phylogeny was a bit disconcerting and calls to question the foundation of separating the genera *Acrasis* and *Pocheina* based on fruiting body morphologies, and the validity of the genus *Pocheina*.

To clarify the taxonomy and systematics of the genus *Pocheina* and the relationship of *Pocheina* spp. to *Acrasis* spp. we collected additional strains of both *P. rosea* and *P. flagellata* from widely separated geographic locales and sequenced their SSU and/or ITS regions (ITS1, 5.8S rRNA gene, ITS2) for comparative analyses. Included in the analyses were newly sequenced ITS regions from all isolates of *Acrasis* spp. studied in Brown et al. 2012, in which only SSU rRNA genes were sequenced. Generation of ITS sequences from these morphologically/phylogenetically delineated *Acrasis* spp. further resolved the relationship between *Acrasis* and *Pocheina* spp. Additionally, these data provided an ideal set of ‘good’ species to assess the benchmark hypotheses generated for *Naegleria* and closely related heteroloboseans, which posited that each species possesses unique ITS sequences and that each genus forms a distinct clade in 5.8S trees (De Jonckheere, 1998; De Jonckheere 2004; De Jonckheere and Brown, 2005). Our results demonstrate that 1) sorocarp morphology correlates with molecular phylogenetic inference, 2) the ‘*P. rosea*’ SSU rDNA sequence reported in Brown et al. (2012) is an obvious contamination from an *A. rosea* isolate, 3) we provide the first publicly available molecular data from *Pocheina* spp. 4) all isolates identified as *Pocheina* spp. form a monophyletic group separate from *Acrasis* spp., which 5) is also monophyletic, and finally, 6) these data are the basis of systematic revisions that establish the monophyly of each major lineage within Acrasidae (including *Allovahlkampfia*).

## MATERIALS & METHODS

### Bark sampling and morphological observation

Bark from *Pinus* spp. trees was collected from about breast height and placed into paper bags from five different sites, including one site (HUNT, yielding isolates HUNT 1 & 2) that was sampled on two separate occasions five years apart (Table 1). The bark was brought into the laboratory, cut into small <1 cm pieces, and placed on sterile weak malt yeast agar (wMY) (0.75 g K_2_HPO_4_, 0.002 g yeast extract, 0.002 g malt extract, 15.0 g agar / liter DI H_2_O) petri plates and hydrated with a drop of sterile DI H_2_O. Plates were incubated at room temperature (ca. 22ºC) under normal ambient light conditions of the laboratory. After 2-7 days, the pieces of bark were scanned for bright pink pocheinoid fruiting bodies using a Leica M205 dissecting microscope (Wetzlar, Germany) with reflected light. Images of fruiting bodies were taken with an attached Canon 650D (Tokyo, Japan) digital camera under reflected light or an Axioskop 2 Plus (Zeiss, Berlin, Germany) at 10X with an attached Canon 650D camera under transmitted light. To observe spore germination, culture slides were created by melting an ∼4 mm x 4 mm block of lactic acid adjusted wMY agar at pH ∼ 5 (as described below) between a slide and cover glass (Brown et al., 2012; Spiegel et al., 2005). After the agar had cooled the cover glass was removed leaving a thin square of solidified agar. A single fruiting body was removed from the bark with a 0.15 mm Austerlitz Insect Pin^®^ (Carolina Biological, Burlington, NC, USA) and placed onto the culture slides along with a drop of DI H_2_O (Brown et al., 2012; Spiegel et al., 2005). Spore germination and trophic cells were observed using an Axioskop 2 Plus compound light microscope equipped with 40X and 63X objectives using both phase contrast and differential interference contrast (DIC) microscopy. Photomicrographs of these cells were taken using a Canon Rebel T2i, Canon 650D, or Canon 5DS digital camera. Culturing attempts of *Pocheina* were made by streaking spores onto wMY agar plates, adjusted to pH ∼5 by adding 3 drops of 5% lactic acid during pouring (Olive et al., 1983), with either: an unidentified species of *Aureobasidium, Rhodotorula mucilaginosa*, or *Escherichia coli*.

**Table 1.**
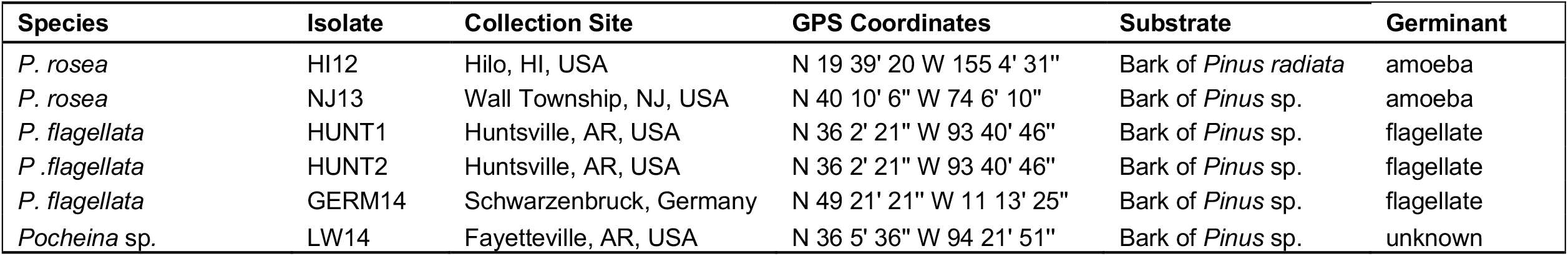
*Pocheina* samples obtained in this study along with their locality, substrate, and the type of cell in which they germinated.

| Species | Isolate | Collection Site | GPS Coordinates | Substrate | Germinant |
| --- | --- | --- | --- | --- | --- |
| *P. rosea* | HI12 | Hilo, HI, USA | N 19 39' 20 W 155 4' 31" | Bark of *Pinus radiata* | amoeba |
| *P. rosea* | NJ13 | Wall Township, NJ, USA | N 40 10' 6" W 74 6' 10" | Bark of *Pinus* sp. | amoeba |
| *P. flagellata* | HUNT1 | Huntsville, AR, USA | N 36 2' 21" W 93 40' 46" | Bark of *Pinus* sp. | flagellate |
| *P. flagellata* | HUNT2 | Huntsville, AR, USA | N 36 2' 21" W 93 40' 46" | Bark of *Pinus* sp. | flagellate |
| *P. flagellata* | GERM14 | Schwarzenbruck, Germany | N 49 21' 21" W 11 13' 25" | Bark of *Pinus* sp. | flagellate |
| *Pocheina* sp. | LW14 | Fayetteville, AR, USA | N 36 5' 36" W 94 21' 51" | Bark of *Pinus* sp. | unknown |

*Allovahlkampfia* (“*Solumitrus*”) *palustris* (PRA325) *sensu* Gao et al., (2022) was purchased from the American Type Culture Collection (ATCC). *Allovahlkampfia* sp. strains BA, OSA and *Al. palustris* were each propagated in either liquid wMY or hay-infusion medium (ATCC 802) in tissue culture flasks supplemented with *E. coli* as the food source.

### Genomic DNA extraction

Genomic DNAs from the *Acrasis* taxa used to amplify the ITS region were from the study of Brown et al., 2012. From new *Pocheina* isolates, two to three sorocarps immediately surrounding the sorocarp taken to observe spore germination were used for DNA extraction. These sorocarps were picked directly from the primary substrate using an ethanol flame-sterilized Austerlitz Insect Pin^®^ and placed directly into 30 μl of Epicentre® QuickExtract™ DNA extraction solution. Aside from the modified volume of solution, DNA was liberated from spores using the manufacture’s recommended protocol. Genomic DNA from *Al. palustris* and strains BA and OSA cell pellets was isolated using the Gentra Puregene Tissue Kit (Qiagen) following the manufacture’s protocol.

### Polymerase Chain Reaction (PCR) from gDNAs

The ITS region (contiguous 3’ end of SSU, ITS1, 5.8S, ITS2, 5’ end of large subunit (LSU) rRNA gene) and SSU rRNA genes were PCR amplified in 25 µl total reaction volumes using Q5^®^ High-Fidelity DNA Polymerase (2x master mix, New England Biolabs, Ipswich, MA, USA) for 30 cycles with combinations of “universal” eukaryotic primers (De Jonckheere and Brown, 2005; Medlin et al., 1988) and custom primers designed against *Allovahlkampfia* spp. and *Acrasis* spp. SSU and ITS sequences (Table 2-4). For each PCR, elongation times were based on slight over-estimates of the expected amplicon size, and annealing temperatures were specified using NEB’s Tm calculator. Post cycling, 20 μl of each PCR reaction was electrophoresed on an 1% agarose gel in TA buffer (9.68 g Tris Base, 2.28 ml glacial acetic acid / liter DI H_2_O) containing SybrSafe (Life Technologies, Grand Island, NY). If weak or no amplicon was seen on the gel, 1 µl of the primary PCR was utilized for nested or semi-nested PCR (Table 2, 3). Upon strong amplification, the DNA bands were cut out of the gel with a razor blade and purified by centrifugation through a 200 µl barrier pipette tip as described in Becker et al. (2024). The ITS region was amplified from two *Allovahlkampfia* strains (BA, OSA), five new *Pocheina* isolates (HUNT1, LW14, NJ13, HI12, and GERM14) as well as from “*Pocheina”* (LOST07L112) and each *Acrasis* spp. from the DNAs isolated by Brown et al., (2012, Table 3). Also amplified were the nearly complete SSU rRNA genes of three new *Pocheina* isolates (HUNT2, NJ13 and HI12) and *Al*.*palustris* (Table 2).

**Table 2.** PCR amplification and product information for the nuclear encoded SSU (18S) rDNA of *Pocheina* and *Solumitrus* amplified for this study.

| Isolate | Species | 1° PCR primers | 2° PCR primers |
| --- | --- | --- | --- |
| HI12 | *P. rosea* | Acd41F : Medlin B | Acd54F : Acd687R, 300F : Acd766R, Acd720F : Acd1424R |
| NJ13 | *P. rosea* | Acd41F : Medlin B | Acd49F : Acd687R, Acd645F : Allo766R, Acd720F : Acd1425R, Acd1380F : Allo1460R, Allo41F : Allo552R |
| Hunt1 | *P. flagellata* | Acd41F : Medlin B | Acd54F : Allo1460R |
| PRA-325 | *Al. palustris* | MedlinA : 1492R |  |

**Table 3.**
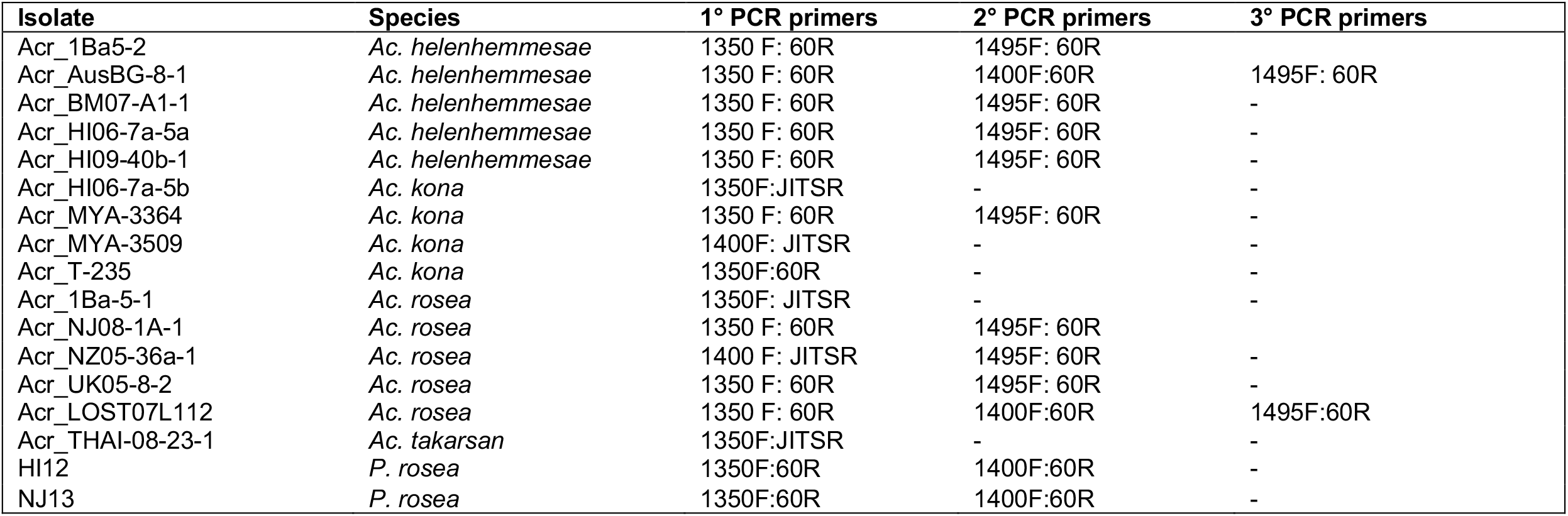
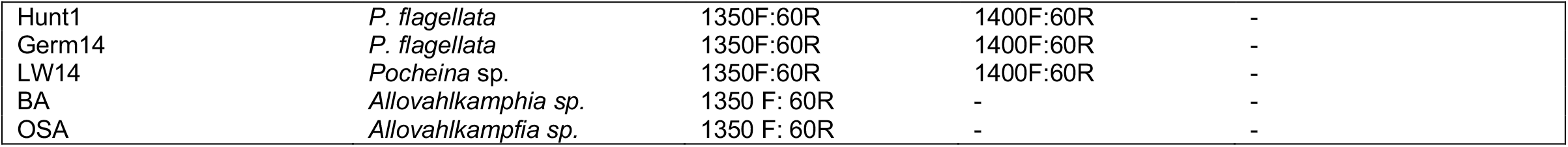
PCR amplification and product information for the ITS1, 5.8S rDNA, and ITS2 regions of all isolates of *Pocheina* and *Acrasis* amplified for this study.

In nearly all instances, PCR products were sequenced directly. In a few cases, weakly amplified amplicons were cloned using the TOPO-Blunt Cloning Kit for Sequencing (Invitrogen, Waltham, MA, USA) following the manufacturer’s protocol. Recombinant plasmids were isolated using the Zyppy Plasmid Miniprep Kit (Zymo Research, Irvine, California, USA) following the manufacturer’s protocol. Samples were Sanger sequenced on an Applied Biosystems 3130xl Genetic Analyzer. Both the SSU rRNA gene and ITS regions were sequenced completely in both directions. All sequences were edited and assembled using Sequencher v. 5.1 (GeneCodes, Ann Arbor, MI, USA). No mixed peaks were seen on the chromatograms for any of our sequences indicating that no microheterogeneity in the SSU rRNA gene or the ITS region exists within or among the cells of any of our isolates.

### Ultra-low input transcriptomics

For *Pocheina* isolate HUNT2, we employed an ultra-low input RNAseq approach to obtain the SSU rRNA and ITS region sequences. A single sorocarp (∼20 cells) was picked from the bark substrate with a 30-gauge platinum wire and placed directly into a 200 µL thin-walled PCR tube. The cells were subjected to a modified version of Smart-Seq2 mRNA extraction and cDNA library preparation (Picelli et al., 2014) that included an additional freeze thaw step for cell lysis, as described in Onsbring et al., (2020). The resulting cDNA library was prepared for sequencing on an Illumina platform using the Nextera XT DNA Library Prep Kit (Illumina, CA, USA) following the manufacturer’s protocol with dual index primers. The library was sequenced using an Illumina HiSeq 4000 at Genome Quebec (Montreal, Canada).

### Transcriptomic assembly and bioinformatic retrieval of the SSU and ITS regions

Low quality bases, adaptor sequences, and Smart-Seq2 primer sites were removed from the HUNT2 raw sequencing read files using TRIMMOMATIC V0.35 (Bolger et al., 2014). The surviving reads were assembled using the de novo assembly program TRINITY V2.1.1 (Grabherr et al., 2011). The full length SSU and ITS region were retrieved using BLASTN and Acrasidae SSU and ITS data to query the transcriptome assembly. The RNAseq data is deposited under the BioProject number XXXXXXXXX.

### Phylogenetic and comparative sequence analyses

Phylogenetic trees were inferred from forty-three SSU rRNA gene sequences, including our new Allovahlkampfiid and *Pocheina* sequences along with other Acrasidae and representative outgroup heteroloboseans (*Naegleria, Willaertia, Pleurostomum*, and *Tulamoeba* spp.). Trees were inferred from an inclusion set of 1,868 unambiguously aligned nucleotide positions using maximum likelihood (ML) and Bayesian inference (BI) methods. Alignments were inferred with MAFFT-LINSI v7.407 (Katoh and Standley, 2013) with default parameters by using the add function; adding new sequences to a seed alignment from Brown et al., 2012. Uninformative sites were removed using BMGE v1.12 (Criscuolo and Gribaldo, 2010) with a maximum gap rate allowed per character set to 0.6 (-g 0.6). A general-time-reversible + gamma distribution (GTR+G) model of nucleotide change was implemented in RAxML v8.2.12 (Stamatakis, 2014) using 25 discrete gamma rate categories. The best scoring ML tree of 300 independent “rapid-hill climbing” tree searches was mapped with topological support assessed by ML analyses of 1,000 nonparametric bootstrap replicates in RAxML under the same model. Bayesian analyses run in MrBayes 3.2.7 (Ronquist and Huelsenbeck, 2012) consisted of two independent Markov chain Monte Carlo (MCMC) runs of 50,000,000 generations printing trees every 1,000 generations with a “burnin” of 7,676,000 generations by which time all parameters converged as assessed by an average standard split deviation (ASD) plateaued at < 0.003 and the potential scale reduction factor convergence diagnostic.

Our new ITS1, 5.8S rRNA gene, and ITS2 sequences of *Pocheina, Acrasis* and allovahlkampfiids were included in pair-wise sequence comparisons, compositional and phylogenetic analyses. Within Heterolobosea, only the 5.8S rRNA gene sequences could be confidently aligned and utilized for phylogenetic analyses. Forty-five 5.8S sequences including those from each new *Acrasis* spp., *Allovahlkampfia* spp., and *Pocheina* spp. along with publicly available 5.8S sequences from other members of Acrasidae and representative outgroup sequences from *Naegleria* spp. were aligned using MAFFT-GINSI with default parameters. Sites not part of the 5.8S and those that were not confidently homologous were removed by hand in Aliview v1.26 (Larrson, 2014). Maximum likelihood and Bayesian trees and support values were inferred as described for the SSU analyses, with the only difference being that the first 8,191,000 generations were discarded as burnin in the Bayesian analysis.

To determine which *Allovahlkampfia* group our new *Allovahlkampfia* BA and OSA isolates belonged to, we conducted unrooted ML and Bayesian analyses of the entire ITS region of all *Allovahlkampfia* strains, as in Gao et al., (2022). All allovahlkampfiid ITS sequences were aligned using MAFFT-LINSI with default parameters. Uninformative sites were removed using BMGE v1.12 with a maximum gap rate allowed per character set to 0.6, resulting in 460bp sites. Maximum likelihood and Bayesian trees and support values were inferred as described for the SSU analyses, with the only difference being that the Bayesian ASD plateaued at < 0.002 and removing the first 6,651,000 generations as burnin.

Finally, we generated a concatenated 5.8S and SSU dataset. To do this we first collected only Acrasidae sequences of both SSU and 5.8S rRNA genes with no outgroup taxa to increase the number of confidently aligned sites. Each gene was aligned using MAFFT-LINSI with default parameters. The SSU alignment was trimmed with BMGE with a maximum gap rate allowed per character set to 0.6. The 5.8S alignment was trimmed by hand. This resulted in 2013 bp from SSU and 170 bp from 5.8S. The SSU and 5.8S sequences were concatenated by hand when the data for both genes was available. In taxa where we only had one of the genes, the missing gene was treated as missing data. This resulted in a dataset of 41 taxa and 2173 nucleotide sites. Maximum likelihood and Bayesian trees and support values were inferred as described for the SSU analyses, with the only difference being that the Bayesian ASD plateaued at < 0.002, and the first 3,716,000 generations were discarded as burnin.

Sequence differences among ITS1 and ITS2 sequences were calculated as uncorrected pairwise differences ignoring gaps between all Acrasidae genera, within each genus, and among species in a genus using the custom script (pdistcalculator.py, https://github.com/socialprotist/pdistcalculator.py/). To do this, alignments of the ITS1 and ITS2 regions were individually analyzed. Sequence repeat regions were assessed by dot blots using YASS genomic similarity search tool (Noe & Kucherov, 2005) comparing each sequence to itself, accessed through the web portal, https://bioinfo.univ-lille.fr/yass/index.php using default parameters.

## RESULTS and DISCUSSION

### Morphological Observations

Five new strains of *Pocheina* were collected for morphological and molecular analyses (Table 1). The morphology of sorocarps and trophic cells of all putative *Pocheina* spp. isolates were characteristic of the genus description (Olive et al., 1983). The fruiting bodies were pinkish orange in reflected light and were made up of a row or rows of wedge-shaped stalk cells topped by a globose mass of spores connected to one another by raised hila (Fig. 1A-G). Slight variation in sorocarp size existed within and among isolates (Fig. 1 A-G). All attempts of *Pocheina* spore germination were successful on wMY agar (adjusted to pH ∼5.0) except for LW14, which consistently failed to germinate. Using the interpretation of Olive et al. (1983), each isolate was assigned to a described species based on the morphology of trophozoites that emerged from spores (Table 1). Isolates were designated *P. flagellata* if a binucleated plasmodium (Fig. 1J) that subsequently cleaved to become 2 uninucleate flagellates emerged from spores (Fig. 1J-L). Isolates were assigned to *P. rosea* if a nonflagellate, uninucleate amoeboid cell emerged from spores (Fig. 1M,N). Though variation was noted among the fruiting bodies seen on *Pinus* bark, we could not predict ahead of time if an amoeba or flagellate would germinate from spores. Germinated trophic cells (either type) did not appear to divide but remained active for 1 h-4 days before the flagellates died, turned to amoebae, or the amoebae encysted. Excystment did not occur under our culturing conditions, even when we passed cysts to fresh agar and food sources. Thus, trophic cells were never again observed after encystment and long-term cultures could not be established for any of the *Pocheina* strains, including the LOST07L112 isolate described in Brown et al., 2012.

**Fig 1.**
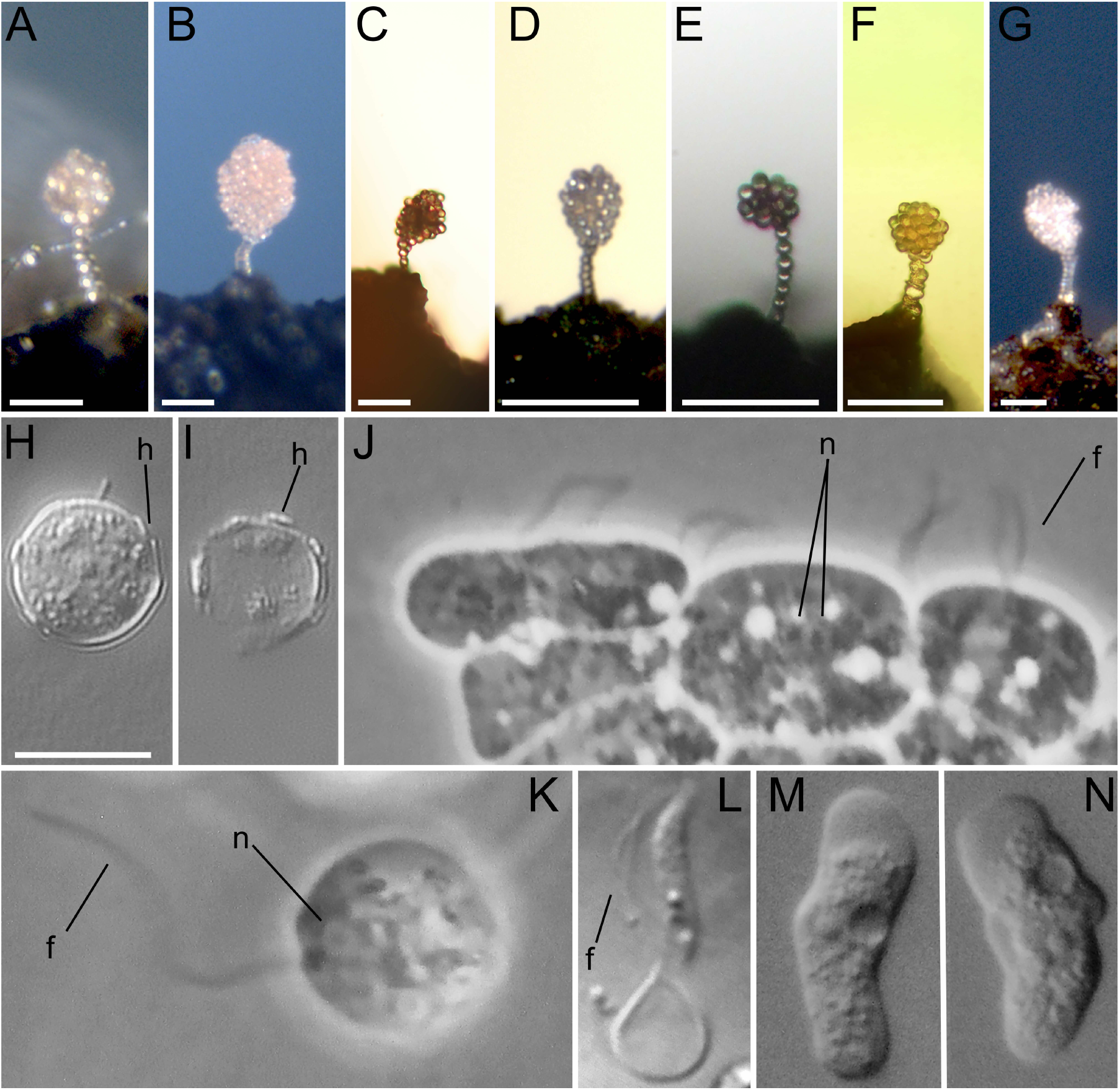
Light microscopy images of *Pocheina* strains. Sorocarps of *Pocheina flagellata* strains GERM14 **(A)**, HUNT1 **(B)**, HUNT2 **(C)**; *Pocheina rosea* isolates HI12 **(D)**, NJ13 **(E)**, LOST07L112 **(F)**; **G)** *Pocheina* sp. LW14. **A-G)** Each scalebar = 50 µm. **A**,**B**,**G** are reflected light and **C-F** are transmitted light. **H)** Spore from *P. rosea* LOST0711L2 with several visible raised hila pointed out (h). **I)** Empty spore wall with raised hila (h). **J)** Germinants from spores from *P. flagellata* strain HUNT1. The center cell is a binucleated amoebae before dividing into a single nucleated flagellate. Two flagella are pointed out in the right most flagellate. **K)** Swimming flagellate from *P. flagellata* HUNT1. **L)** Elongated flagellate from *P. flagellata* GERM14, flagella (f). **M, N)** Amoebae from *P. rosea* isolate NJ13. Image H-N are to scale, scalebar = 10 µm.

The amoebae of *P. rosea* (isolates HI12 and NJ13) moved with eruptive pseudopodia, as did the amoebae that emerged from germinated spores of the LOST07L112 isolate (Brown et al., 2012). *P. flagellata* cells (isolates HUNT1, HUNT2, GERM14) were semi-amoeboid when emerging from spores (Video S1). Once the morphology of flagellated trophozoites stabilized, they had a constant body shape and swam using their two anterior flagella (Fig. 1K,L, Video S1-2). As noted by Olive et al. (1983), flagellate morphology was variable among isolates, which parallel our observations of the HUNT and GERM isolates. The flagellated cells of HUNT (1 & 2) were spherical to obovate in shape with a short yet distinct rostrum (Fig. 1J). The flagellated cells of GERM14 were narrow, elongated, and tapered at the posterior end (Fig. 1L). On agar culture slides of HUNT1, which were kept for several days, flagellates transitioned into crawling nonflagellate amoeboid cells, as previously reported for *P. flagellata* (Olive et al., 1983). We did not directly observe the transition in real time, but only saw that amoebae were present on older agar culture slides. This flagellate to amoeba transformation is associated with the vast majority of heteroloboseans (Panek et al., 2016) and may be an ancestral trait for the entire lineage. Although the agar culture slides made photo-documentation difficult, videos captured the morphological essence of flagellated cells (Video S1-2).

### Phylogenetic and Molecular Results

The nearly complete SSU rRNA gene for *P. rosea* isolates HI12 and NJ13 and *Al. palustris* were generated using a PCR approach, while the SSU rRNA sequence from *P. flagellata* HUNT2 was bioinformatically recovered from its transcriptome, as well as the ITS region. Concurrently we generated the complete nuclear encoded ITS region (with the 3’ end of the SSU, ITS1, 5.8S, ITS2, 5’ end of the large subunit rRNA gene) for six *Pocheina* isolates (5 new strains plus LOST07L112), each of the *Acrasis* spp. isolate from Brown et al. 2012, and two *Allovahlkampfia* spp. through PCR amplification.

Many Acrasidae and other heterolobosean taxa have group I introns within their SSU rRNA genes (Brown et al., 2012), including two of our new *P. rosea* isolates (HI12 and NJ13, Fig. 5). The naming system of group I introns corresponds to their position in the *E. coli* SSU (16S) rRNA gene (Johansen and Haugen, 2001; GenBank accession AB035922). *Pocheina rosea* isolate HI12 minimally possesses 3 group I introns that are located at sites known to be common intron sites (S516, S895, and S1199). These introns range from 264bp to 1026bp. We cannot be certain if this isolate’s SSU has additional introns because we were not able to obtain its complete SSU rRNA gene and are missing a site that commonly possesses group I introns in other Heterolobosea (i.e., the gap in HI12’s SSU sequence encompasses the 18 bp before and 11 bp after the S956 group I insertion site, Fig. 5). *Pocheina rosea* NJ13 has more group I introns (five) in its SSU rRNA gene than any other published Acrasidae SSU; there is one at each known Acrasidae insertion site (S516, S895, S956, S1199), as well as a novel insertion site (S1211) not yet observed in Acrasidae. Two of the introns, S516 and S956, have an embedded open reading frame (ORF) encoding a putative 189aa His-Cys box homing endonuclease (HEG). The S516 intron’s HEG is in the forward direction in frame +2 at 14bp 3’ of the intron’s insertion site. The S956 intron’s HEG is in reverse direction in frame -2 starting at 220bp from the 3’ end of the 1081bp intron (Fig. 5). These two predicted proteins are not easily aligned with one another and share limited homology, with only a few short stretches (ca. 40 aa) of 30-40% amino acid identity. The S516 HEG of NJ13 shares 60% amino acid identity to that of the S516 HEG in *Acrasis rosea* 1Ba5-1 (GenBank AER08052). The S956 HEG protein blasts (BlastP) to a HEG protein found on a short contig of the genome of *Ac. kona* strain ATCC MYA-3509 (JAOPGA020000628.1) (Sheikh et al., 2023). This 2114bp *Ac. k*ona genomic contig blasts (BlastN) to the SSU rRNA gene (HM114344) of the same strain but only shares a short 72 bp homologous stretch with just 81% nucleotide sequence identity. The contig itself does not appear to be a rRNA gene, only having this short stretch of the SSU rRNA gene and the coding sequence for a HEG. Introns S516 of *P. rosea* HI12 and NJ13 are nearly the same length (1026-1027bp) and the last 874bp of the intron is nearly identical. However, the first 152bp, including where the start codon a functional HEG would be (if present), are dissimilar and not alignable. Contrary to strain NJ13, no in-frame start codon is present in strain HI12 and all conceptual translations lead to frameshifts and no obvious HEG ORFs. Future investigation of the patterns of introns and HEGs is necessary to tell the full story of intron evolution of the clade.

The topology of the SSU rRNA gene phylogeny (Fig. 2) shows that all newly isolated *Pocheina* spp. form a fully supported clade with 100% Maximum Likelihood bootstrap support (BS) and Bayesian posterior probability (PP) of 1.0 within a fully supported (100%/1.0) Acrasidae *sensu* Brown et al. (2012). Both *P. rosea* (NJ13 and HI12) isolates branch together with high support (100%/1.0) and are sister to the single *P. flagellata* (HUNT2) SSU rRNA sequence. However, the sequence identified as “*P. rosea”* LOST07L112 in Brown et al. (2012) remains problematic because it branches within the genus *Acrasis* in a fully supported clade of the species *Ac. rosea*. We will later demonstrate that this sequence is a contaminant from a verified *Ac. rosea* isolate, and as such, the genus *Acrasis* is likewise a highly supported monophyletic group in SSU trees, (98%/1.0) and now completely conforms to the species concept of Brown et al. (2012) without exceptions. A clade with moderate BS and full PP support containing soil amoeba AND12 (AY965862) and all *Allovahlkampfia* spp. is recovered (71%/1.0). This is congruent with the results of Geisen et al. (2015), although they recovered higher ML support for the clade (possibly because of differing taxa in their analyses). Overall, three monophyletic lineages are resolved in SSU analyses of Acrasidae: *Acrasis, Pocheina*, and *Allovahlkampfia*. There is limited resolution among the three major Acrasidae lineages, but the SSU tree shows a basal split between *Acrasis* and a poorly supported group comprising *Pocheina* as sister to *Allovahlkampfia* spp. (65%/0.92).

**Fig 2.**
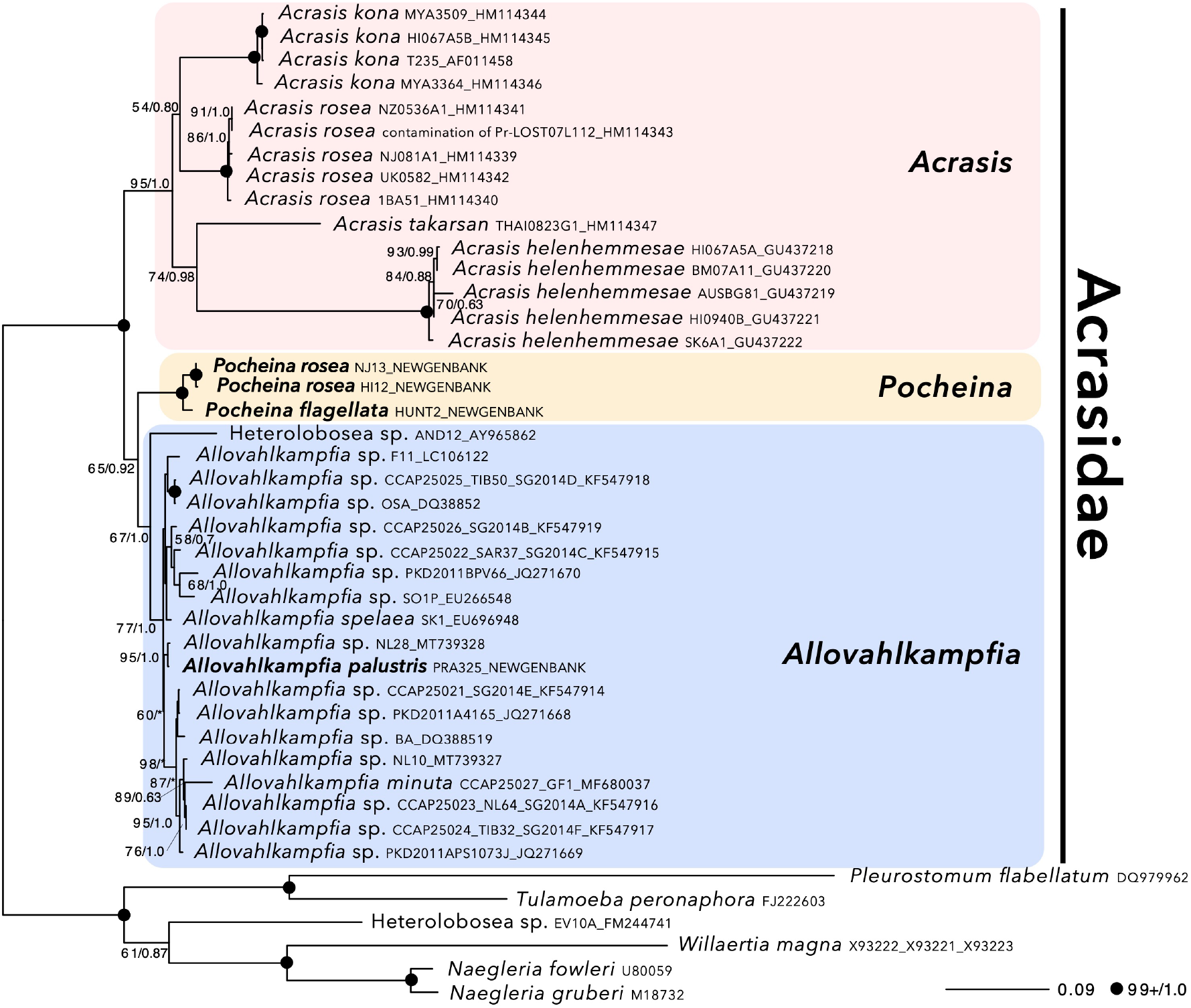
Maximum likelihood SSU rRNA gene tree of Acrasidae with closely related heteroloboseans as an outgroup, using RAxML with GTR+G model of substitution. Bootstrap support values and Bayesian posterior probabilities (see Methods) are shown at nodes with support values above 50%/0.7, respectively. An asterisk (*) denotes PP below 0.5. Our novel data are bolded.

The newly sequenced ITS region of all *Pocheina* isolates, the *Acrasis* isolates from Brown et al. (2012), and *Allovahlkampfia* isolates OSA and BA, were used to test the taxonomic concepts advocating that genera are monophyletic in 5.8S phylogenetic trees and species can be delineated by unique ITS1/2 sequences (De Johnkheere, 2004; De Johnkheere and Brown, 2005). We took advantage of the fact that *Acrasis* spp. comprise ‘good’ species that are readily identifiable using a combination of morphological and SSU molecular data (Brown et al., 2012). The ITS sequence data was instrumental in honing the systematics of taxa comprising each of the major Acrasidae lineages. These data codify the discrepancy of the morphology of ‘*Pocheina*’ LOST07L112 and its position within SSU phylogenetic trees, provides further support for the monophyly of *Allovahlkampfia*, and establishes the monophyly of the genus *Pocheina*. In addition, these data highlight the need to be cautious when naming new species based solely on molecular data.

Because the 5.8S rRNA gene is short and highly conserved, relationships among the major lineages included in its phylogenetic analyses are weakly supported (or unresolved) and not appropriate to infer broad-scale relationships (Fig. 3). However, the 5.8S phylogeny supports most of the salient interpretations inferred from the SSU rRNA gene analyses. In the 5.8S analyses, Acrasidae is recovered as a fully supported clade (100%/1.0), though the genus *Acrasis* appears paraphyletic. Consistent with the SSU phylogeny, each species of *Acrasis* is recovered with strong support. Most notable is that with greater taxon sampling, the genus *Pocheina*, including ‘*P. rosea*’ LOST07L112 is fully supported (100%/1.0) (and weakly sister to *Allovahlkampfia*).

**Fig 3.**
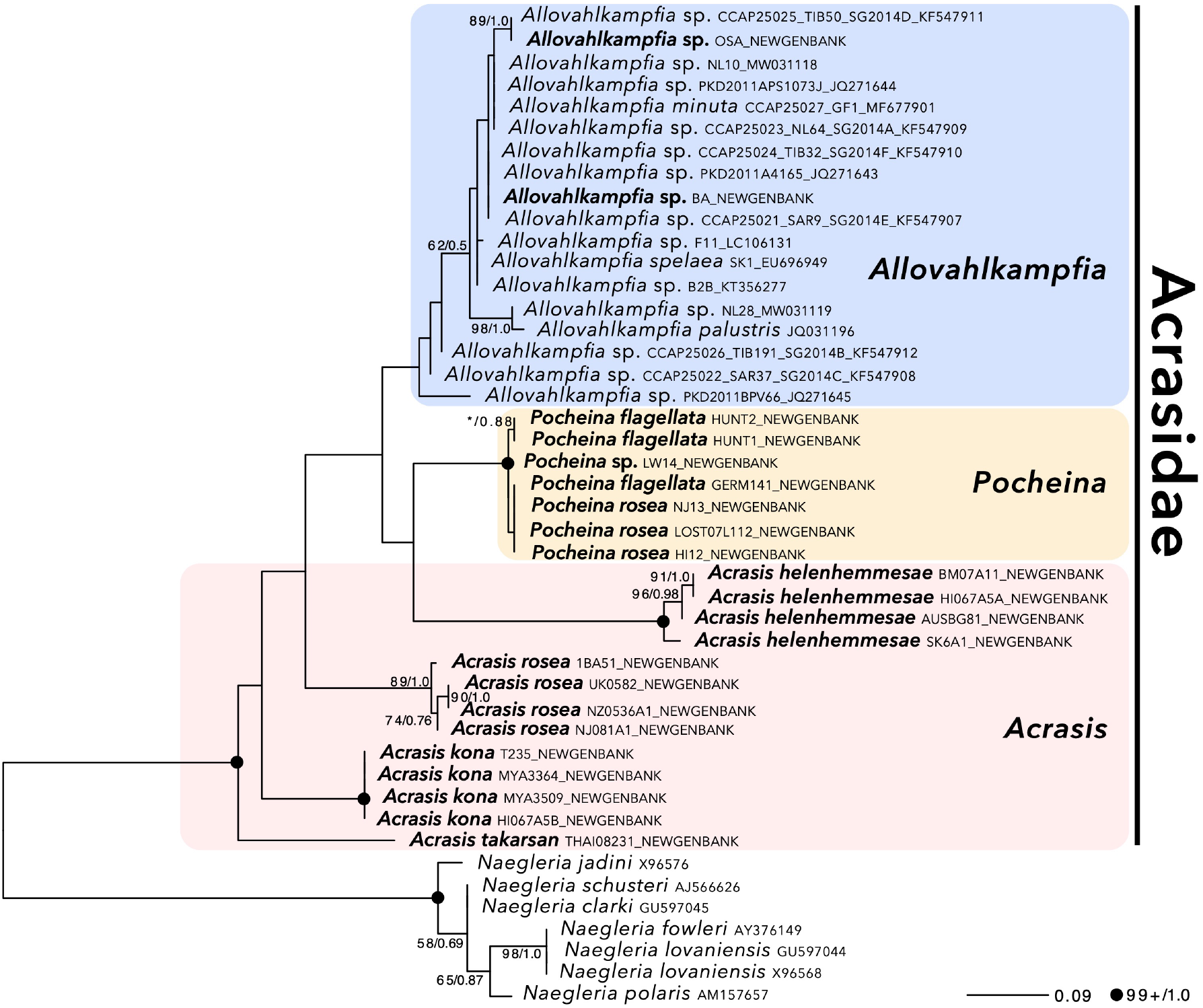
Maximum likelihood 5.8S rRNA gene tree of Acrasidae with closely related heteroloboseans as an outgroup, using RAxML with GTR+G model of substitution. Bootstrap support values and Bayesian posterior probabilities (see Methods) are shown at nodes where both values are above 50% or 0.5, respectively. The asterisk (*) denotes BS below 50%. Our novel data are bolded.

The strongly supported and conflicting position of ‘*P. rosea*’ LOST07L112 sequences in their respective SSU and 5.8S rRNA gene phylogenetic trees is problematic (Fig. 2, 3). The SSU sequence is fully supported as a member of *Ac. rosea* while the 5.8S sequence is fully supported as a member of the genus *Pocheina*. Fortunately, analyses of the independently generated SSU and ITS regions from all *Pocheina* and *Acrasis* isolates provide a logical resolution to this phylogenetic inconsistency.

When the ‘*P. rosea*’ LOST07L112’ SSU sequence was generated (Brown et al. 2012), there were no other molecular data from other *Pocheina* strains for comparisons. Sequence comparisons became possible only when we generated molecular data from new *Pocheina* isolates, particularly the ITS region from each of our *Pocheina* spp. and *Acrasis* spp. strains, which overlap to varying extents with the independently amplified SSU rRNA genes reported in Brown et al. (2012). The first piece of evidence that the ‘*P. rosea*’ LOST07L112 ITS1-5.8S-ITS2 and SSU (HM114343) sequences do not originate from the same organism is that there are 18 nucleotide differences among the overlapping 105 bp SSU rRNA gene(s) shared between the separately amplified ITS and SSU rRNA regions (Fig. 4), even though the exact same genomic DNA was used to assemble the amplification reactions. This contrasts with the minimal intra-strain sequence differences of this region among all *Ac. rosea* isolates generated by SSU rRNA gene amplifications (0-2 bp), and the 100% SSU sequence identity amongst all the *Pocheina* spp. ITS amplicon and transcriptome generated sequences. The pairwise sequence difference between the SSU and the ITS amplicons of ‘*P. rosea*’ LOST07L112 is well outside this range.

**Fig 4.**
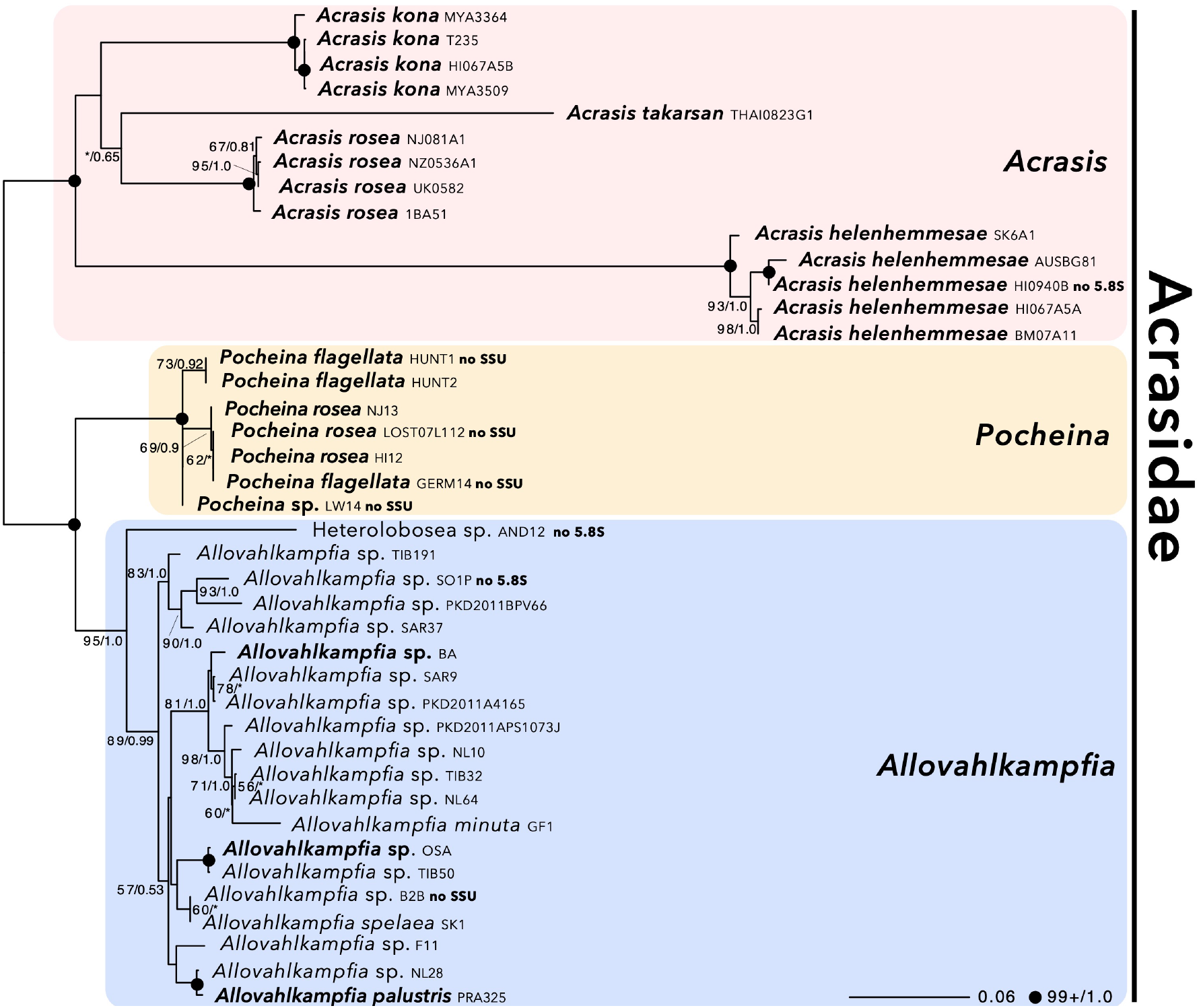
Maximum likelihood tree of Acrasidae SSU rRNA and 5.8S rRNA genes concatenated, using RAxML with GTR+G model of substitution. Bootstrap support values and Bayesian posterior probabilities (see Methods) are shown at nodes where both values are above 50% or 0.5, respectively. An asterisk (*) denotes BS or BI below 50% or 0.5, respectfully. Taxa in which a gene is missing is denoted as “no SSU” or “no 5.8S”. Taxa for which novel data are presented here are bolded.

**Fig 5.**
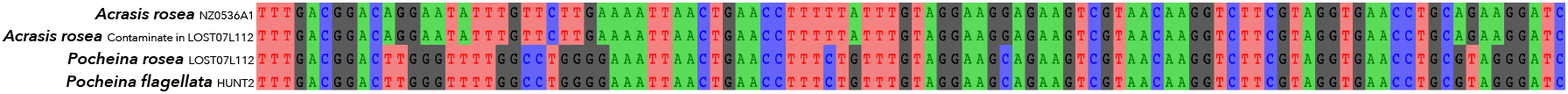
Image of an alignment of the 3’ end of the SSU rRNA gene from LOST07L112 from Brown et al. 2012 (HM114343) and from the ITS region amplification obtained here. The top line is the sequence from *Acrasis rosea* NS05-36a-1 (HM114341). The bottom line is from *Pocheina flagellata* HUNT2 obtained here.

**Fig 6.**
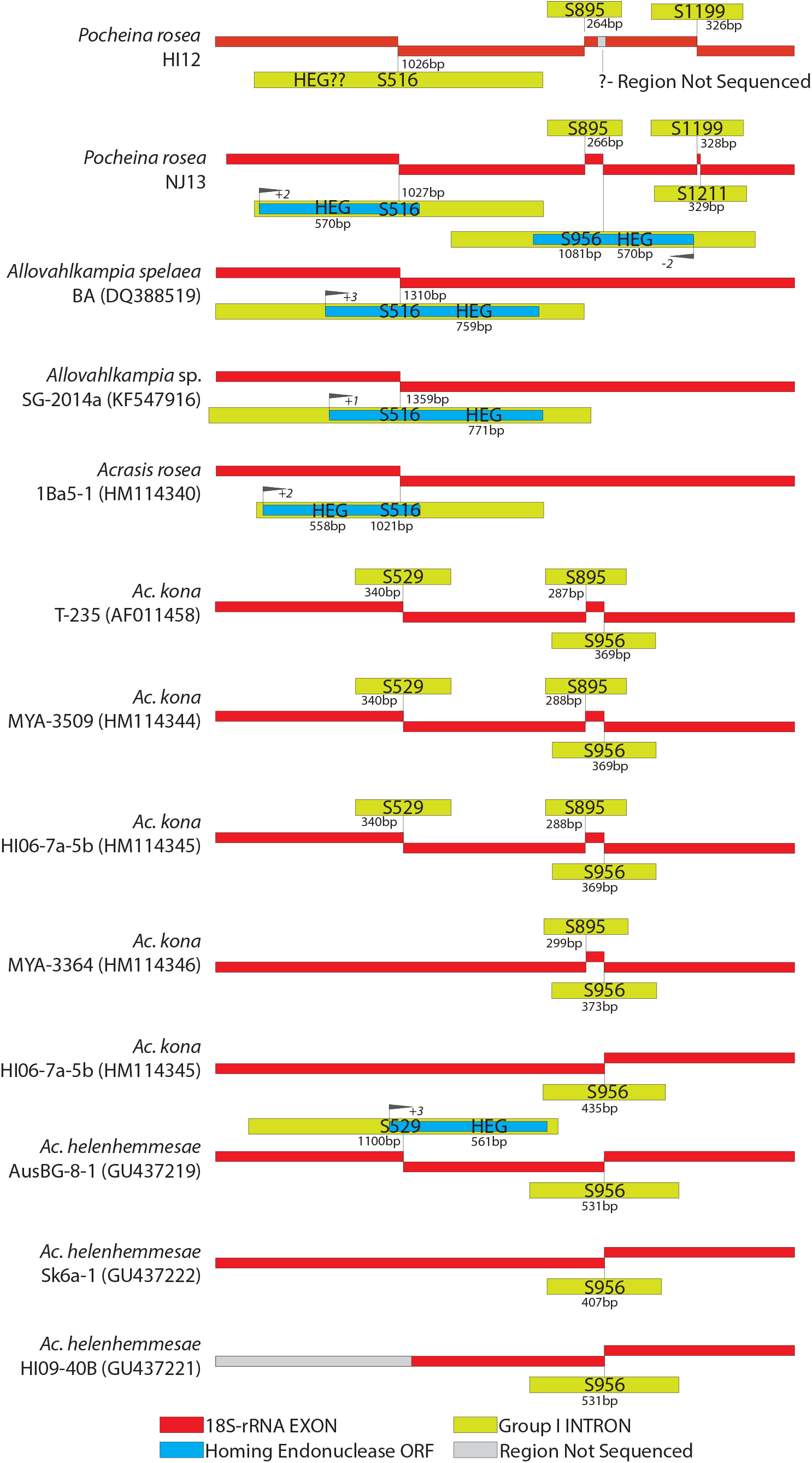
Map of known group I introns and homing endonucleases in Acrasidae nuclear encoded SSU rRNA genes. The location of each intron is depicted with SNNN, representing the homologous base in the 16S rRNA gene of *E. coli*. Red lines are rRNA coding regions. Yellow lines are group I introns. Blue boxes within group I introns are homing endonuclease open reading frames (ORF). Grey lines are regions that were not sequenced. All lines are to scale.

We then determined which gene sequence belongs to *Pocheina* and which to the contaminant. Because all our newly generated ITS sequences are contiguous with the SSU rRNA gene, we were able to link each ITS to its corresponding SSU rRNA gene. This was not possible with SSU rRNA gene sequences amplified in the Brown et al. (2012) study because the 3’ reverse PCR primers were within the SSU rRNA gene. The nearly complete SSU and ITS sequences of *P. flagellata* (HUNT2)/*P. rosea* (NJ13, HI12) can each be assembled into a contiguous contig with 100% sequence identity in the overlapping SSU rRNA gene and the SSU and 5.8S phylogenic tree topologies are congruent. The same is true for all *Acrasis* spp. isolates except for ‘*P. rosea*’ LOST07L112. Besides sorocarp morphology and the 100% SSU sequence identity amongst all contiguous *Pocheina* ITS region sequences (discussed above), the remainder of the LOST07L112 ITS can be fully aligned with those from all other *Pocheina* isolates and lacks similarity to the ITS1/ITS2 of any *Acrasis* isolate. Thus, multiple lines of evidence indicate that the ITS region sequence of ‘*P. rosea*’ LOST07L112 originated from *Pocheina*.

On the other hand, we can confidently assign the SSU rRNA gene sequence attributed to ‘*P. rosea*’ LOST07L112 (HM114343) to an *Ac. rosea* contamination. There is no branch-length between HM114343 and *Ac. rosea* NZ0536A1 (HM114341) in SSU phylogenetic analyses (Fig. 2) and close examination of their edited SSU rRNA gene sequences reveal that they are 100% compatible with one another. These sequences differ in only 8 positions where mixed peaks on the sequencing chromatograms of one was fully resolved to a compatible base in the other (Table 4 of Brown et al., 2012). The amplicon yielding HM114343 likely originated from a pipetting error of *Ac. rosea* NZ0536A1 DNA into the LOST07L112 PCR tube during assembly of the SSU amplification reaction. This is very plausible because the PCRs from Brown et al. (2012) were set up in concurrent experiments. Thus, we are reassigning the ‘*P. rosea’* LOST07L112 SSU sequence to *Ac. rosea* LOST07L112 (HM114343).

**Table 4.**
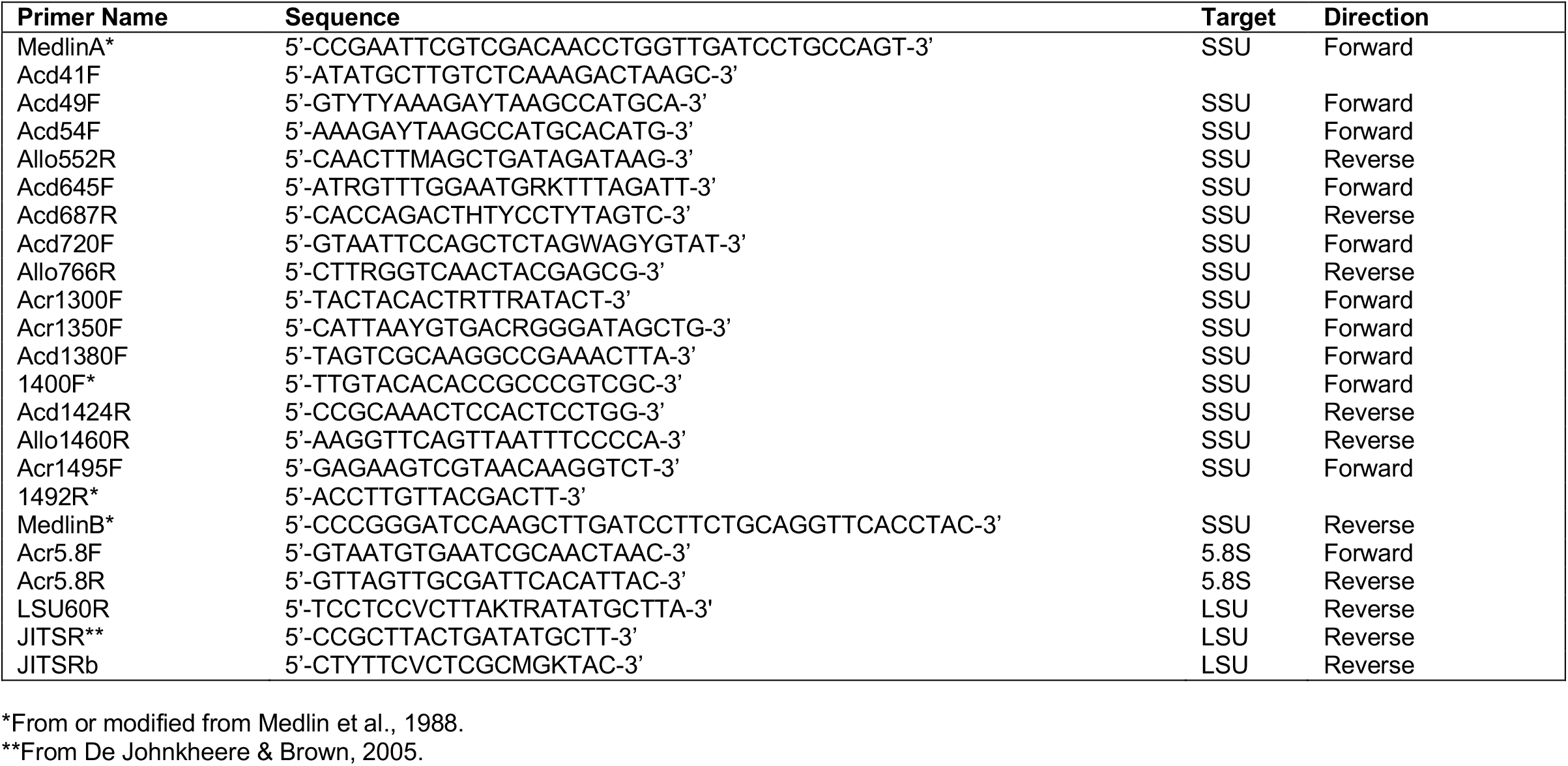
Primer names and sequences used for the amplification of the nuclear encoded SSU (SSU) rDNA, ITS1, 5.8S rDNA, and ITS2 of all isolates of *Pocheina, lumitrus* and *Allovahlkampfia* amplified for this study.

### Systematics and Taxonomy of Acrasidae

Based on fruiting body morphology, members of the genus *Acrasis* and of the genus *Pocheina* are readily distinguishable on the primary isolation substrates, which thus far comprise plant materials such as bark, leaves, or inflorescences (Olive et al., 1983; Brown et al., 2010, 2012). Members of *Allovahlkampfia* are currently circumscribed by rRNA sequence data and the clade comprising the genus is only moderately supported in single gene phylogenies (Fig. 2, 3) although there is increased support in concatenated SSU+5.8S trees (Fig. 4; Gao et al., 2020). Even though there are some morphological differences among the three described species of *Allovahlkampfia* (Anderson et al., 2011; Del Valle and Maciver, 2017; Walochnik and Mulec, 2009), it is unclear if they are taxonomically informative. The ITS-based species concept (De Jonckheere 2004) for *Allovahlkampfia* is not possible to address now. However, *Allovahlkampia* was annotated into 5 groups based on unrooted phylogenetic analysis based on the entire ITS region (Gao et al., 2022). The newly sequenced ITS region of *Allovahlkampfia* strains BA and OSA branch with group 2 and group 3, respectfully (Fig. S1). Molecular phylogenetic analyses containing more genes should better delineate *Allovahlkampfia*, which could lead to the recognition of taxonomically informative characteristics, as was the case with *Acrasis* (Brown et al., 2012).

The 5.8S rRNA gene delineation of genera is not especially useful in Acrasidae because of its limited resolution in conjunction with the long branch leading to the outgroup taxa (Fig. 3). However, 5.8S phylogeny does recover the genus *Pocheina* with full support (Fig. 3) and concatenation of 5.8S and SSU rRNA gene sequences and even SSU alone seems to have enough information to delineate genera amongst the Acrasidae (Fig. 2, 4).

We have multiple isolates of ‘good’ species within *Acrasis* to assess whether each species possess unique ITS sequences. Considering only ITS sequences (not including the 5.8S rRNA gene), in most species for which we have more than one isolate, there are intra-specific sequence differences (Table S1). Except for *Ac. kona* strains which have identical ITS1 and ITS2, no *Acrasis* species has a completely unique ITS shared exclusively amongst strains. The same is true for *Pocheina*, as the only strain sharing identical ITS sequences are HUNT 1 & 2 (which are likely the same strain; see below), collected from the same tree years apart. The most extreme example of intra-specific ITS sequence diversity is from the morphologically simplest *Acrasis, Ac. helenhemmesae* (Brown et al., 2010). This species has the longest ITS of all currently known *Acrasis* species (Table S1). Among the four *Ac. helenhemmesae* isolates, the ITS regions (ITS1 and ITS2, excluding the 5.8S rRNA coding gene) ranges from 3094 to 4275 nucleotides in length. Not only are there length differences, but scattered between regions of sequence similarity, there are multiple regions that are unalignable. Much of this can be attributed to numerous direct and indirect repeats within the ITS, which also account for some of the intra-specific ITS sequence length differences (Fig. S3). Thus, the hypothesis of species delineation based on unique ITS sequences (De Jonckheere, 2004; De Jonckheere and Brown, 2005) does not hold for *Acrasis*, and also, may not be applicable for *Pocheina*. We can only urge caution when establishing molecular barcodes in the absence of independently verifiable taxonomic criteria.

Incorporating molecular data into a species concept for members of the genus *Pocheina* is in its infancy. Currently, there is a paucity of isolates to study, and it seems to be relatively rare in the environment, as it is rarely observed or reported. We simply have too few isolates to confidently determine the taxonomic significance or stability of morphological variations observed in any life-history stage (this study; Olive et al., 1983), especially since isolates could not be cultured, and thus, not amenable for growth in a “common garden” environment. Replicates are required to assess the support and stability of an observation. Our only example comes from *P. flagellata* HUNT 1 & 2, which were recovered from the same tree, 5 years apart. Germinating trophozoites possessed the same morphologies and are nearly identical at the molecular level. We interpret this to mean that the tree was colonized by this strain, which possesses morphological characteristics that are stable over time. Additional replicates of other isolates are warranted test this supposition.

Taxonomic designations are hypotheses that are subject to re-interpretation when additional data become available. We do not have enough information yet to challenge the taxonomic definitions of *P. flagellata* vs *P. rosea* (Olive et al., 1983). However, detailed analyses of the ITS region suggest that their taxonomy may be subjected to revisions. The 5.8S phylogenetic tree shows *P. flagellata* GERM branching with the *P. rosea* isolates rather than the other *P. flagellata* isolates (HUNT), albeit with very poor support (Fig 3). Close inspection of the ITS region alignment hints at ‘signature sequences’ shared between *P. flagellata* GERM and *P. rosea* to the exclusion of *P. flagellata* HUNT (Fig. S2). It may turn out that the differing morphology among flagellates becomes a component in splitting *P. flagellata* into different species. Unfortunately, Olive et al. (1983) illustrates morphologically unique flagellates from multiple strains in their image plates, but the image of the type-strain (NC81-87) is not denoted. Thus, interpretation of the morphological characteristics of the type-strain of *P. flagellata* Olive et al. (1983) is not possible. More isolates, more data and additional analyses are needed prior to any taxonomic revision within *Pocheina*. Unfortunately, our attempts to generate a stable culture of *Pocheina* have failed. Without such, generating conclusive morphometric data is not possible or practical.

### Ancestral traits of Acrasidae

To date, nearly all *Allovahlkampfia* species have been isolated from soil environments and typically cultivated as amoebae in liquid media, with occasional cultivation on agar plates (Anderson et al., 2011; Del Valle and Maciver, 2017; Gao et al., 2022; Geisen et al., 2015). None have been isolated as a fruiting amoeba, unlike *Acrasis* and *Pocheina* that have exclusively been isolated from fruiting bodies from plant materials. To date, the only Allovahlkampfia to be isolated from plant material is strain BA. It was originally isolated from tree bark as an amoeba and propagated in this form. However, a single sorocarp was induced in a study by Brown et al. (2012) that displayed a morphology distinct from both *Acrasis* and *Pocheina;* notably lacking the raised hila on spores. Induction of the sorocarp was achieved by adding amoebae to *Pinus* sp. bark soaked in a water/yeast slurry (Brown et al., 2012). In that same study, *Allovahlkampfia* sp. strain OSA, which was isolated from the fruiting body of a basidiomycete jelly fungus *Dacrymyces* sp., failed to undergo fruiting in similar attempts (Brown et al., 2012).

We are unaware of efforts to induce fruiting body formation in other *Allovahlkampfia* strains. Thus, it is plausible that some *Allovahlkampfia* are sorocarpic amoebae and that appropriate conditions for inducing cell aggregation and/or fruiting body formation have yet to be discovered. Based on our phylogenetic analyses, it is most parsimonious to propose that social multicellularity and fruiting body formation are ancestral traits in the Acrasidae lineage, as fruiting is observed across major clades. It is possible that some isolates, strains, or species have lost the ability to form fruiting bodies through evolution, or that this ability has simply not been observed under laboratory conditions. Therefore, the absence of fruiting should not be considered a taxonomically significant feature.

It is conceivable that *Allovahlkampfia* (and *Pocheina*) may eventually conform to a morphological and molecular species concept similar to that used for *Acrasis*. Future efforts to induce cell aggregation and fruiting body formation would be valuable for comparative morphological studies, taxonomic assignments based on multiple independent traits, and gene expression analyses that could elucidate the similarities and differences in cellular aggregation (Sheikh et al., 2023). In a similar vein, given the presence of flagellated cells in *Pocheina* and one undocumented observation in *Acrasis* (see Brown et al., 2012, pp 104), testing for flagellates throughout the lineage could further uncover their occurrence where they are currently undocumented.

## Conclusion

Sorocarpic amoebae occur across the eukaryotic tree and are found in all major lineages containing amoeboid taxa. The ability to form fruiting bodies mediated by cell-cell aggregation likely evolved independently at least eight times (Tice and Brown, 2022). Most of these lineages are mono-typic genera or, at best, contain just a few species. Examples include *Copromyxa protea* (Tubulinea, Amoebozoa), *Fonticula alba* (Holomycota, Obazoa), *Guttulinopsis* spp. (Cercozoa, Rhizaria) and *Sorodiplophrys stercorea* (Labyrinthulomycetes, Stramenopiles) (Brown et al., 2009; 2010; 2012; Brown and Silberman, 2013; Raper et al., 1977; Schuler et al., 2018; Tice et al., 2016; Tice and Brown, 2022). However, sorocarpic amoebae are far from obscure protists. The most famous example may be *Dicyostelium discoideum* (Evosea, Amoebozoa); it is a model organism for the study of cell motility, chemotaxis, pattern formation, host-pathogen interactions and numerous biomedical processes (Bozzaro, 2019; Martin-González, et al., 2021). The dictyostelids are speciose and highly successful in soil environments (Sheikh et al., 2018). So far, Acrasidae is the only other group containing a rich diversity of sorocarpic amoebae; comprising three genera and multiple sorocarpic species. Complementing ever-improving traditional culturing and molecular methods for detecting biodiversity, expanded global sampling into underexplored environments (e.g., dead, decaying, or living plant material) is likely to uncover additional acrasid species, and perhaps even genera. It is likely that such studies may also reveal additional diversity in other lineages of sorocarpic amoeba, which would provide a wealth of taxa amenable to comparative studies and perhaps even model organism development.

## DATA AVAILABILITY

The SSU rRNA gene sequences and the raw Illumina sequencing reads have been deposited on NCBI under XXXXXXXXX and in Bioproject XXXXX respectively. The transcriptome assembly from HUNT2 and untrimmed and trimmed SSU and ITS alignments are available on figshare, https://doi.org/10.6084/m9.figshare.27174660.

## ACKNOWLEDGEMENTS

This work was supported by the Arkansas Bioscience Institute awarded to JDS and the United States National Science Foundation (NSF) Division of Environmental Biology (DEB) grant 2100888 (http://www.nsf.gov) awarded to MWB. We thank Dr. Don Hemmes and Miss Lucy C. Silberman for their effort in collecting substrates from Hawaii and Germany, respectively. Preliminary work on this project was also supported by National Science Foundation Grants DEB 0329102 and DEB 0316284 awarded in part to FWS.

## SUPPLEMENTAL DATA

**Fig S1.**
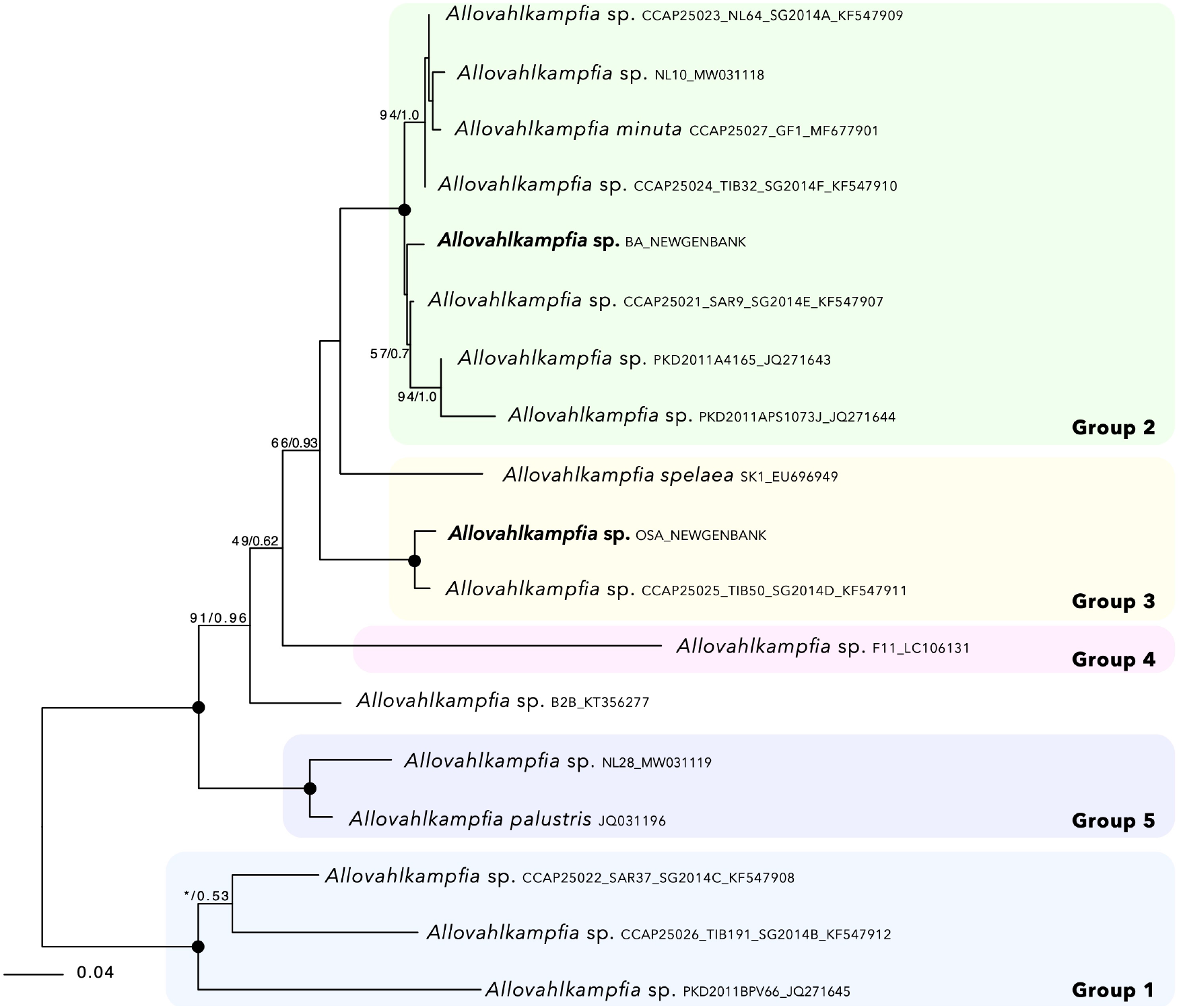
Maximum likelihood tree of the whole ITS region of *Allovahlkampfia* spp. using RAxML with GTR+G model of substitution. Groups of Gao et al., are denoted. Bootstrap support values and Bayesian posterior probabilities (see Methods) are shown at nodes where both values are above 50% or 0.5, respectively. The asterisk (*) denotes BI below 0.5. Nodes with full BS/PP support are denoted with solid circles. Taxa for which novel data is presented here are bolded.

**Fig. S2.**
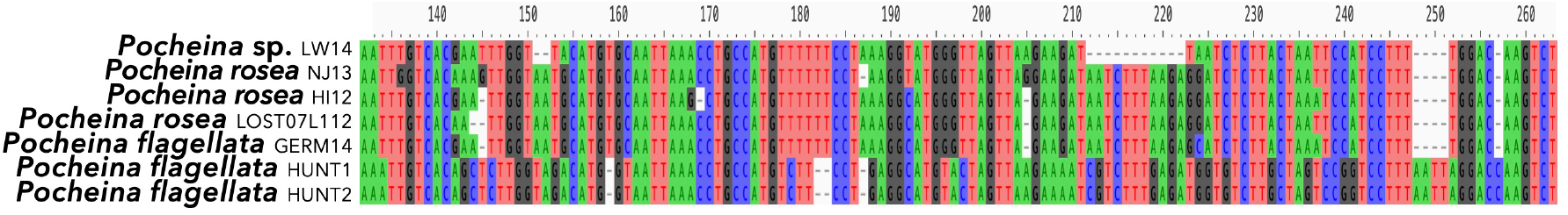
Image of an alignment of a 133 bp region of ITS1 gene from *Pocheina* spp. illustrating the variability in the ITS1 between the HUNT strains (1 & 2) and (bottom two lines) compared to the other *Pocheina* strains as well as the similarity of *P. flagellat*a GERM15 to *P. rosea* sequences.

**Fig. S3.**
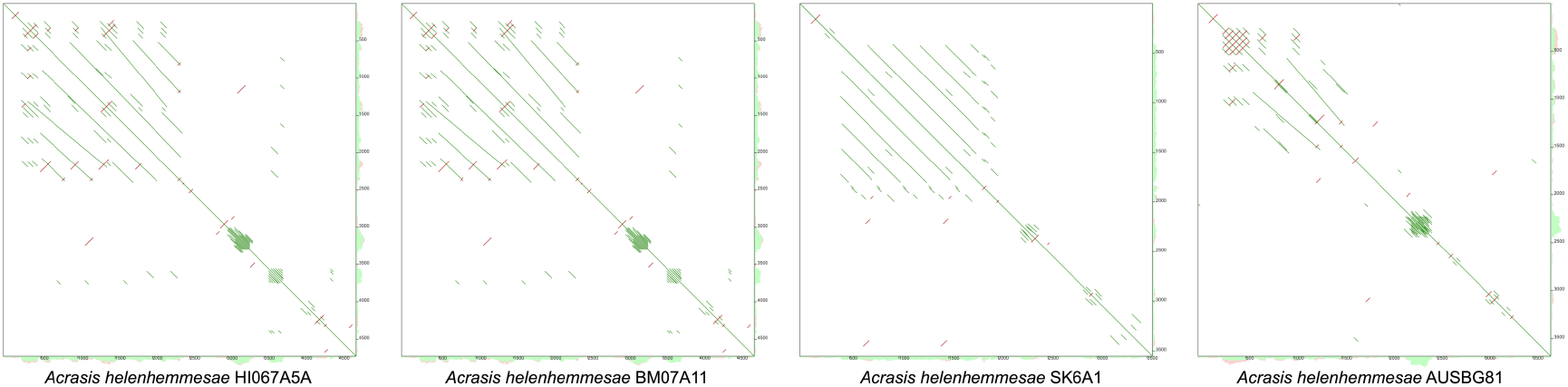
Dot blots inferred of the ITS1 for each strain of *Acrasis helenhemmesae*. Inferred with YASS genomic similarity search tool by comparing each sequence to itself. Direct repeats are in green off the center line and inverted repeats are in red.

**Video S1**. Video microscopy of *Pocheina flagellata* isolate HUNT1. Video is in real time.

**Video S2**. Video microscopy of *Pocheina flagellata* isolate GERM14. Video is in real time.

**Supplemental Table S1**. Uncorrected pairwise distance matrix of aligned ITS1 and ITS2 sequences from Acrasidae. Table also includes the length of the ITS sequences.

